# *In vivo* corneal and lenticular microscopy with asymmetric fundus retroillumination

**DOI:** 10.1101/2020.03.10.985341

**Authors:** Timothy D. Weber, Jerome Mertz

## Abstract

We describe a new technique for non-contact *in vivo* corneal and lenticular microscopy. It is based on fundus retro-reflection and back-illumination of the crystalline lens and cornea. To enhance phase-gradient contrast, we apply asymmetric illumination by illuminating one side of the fundus. The technique produces micron-scale lateral resolution across a 1-mm diagonal field of view. We show representative images of the epithelium, the subbasal nerve plexus, large stromal nerves, dendritic immune cells, endothelial nuclei, and the anterior crystalline lens, demonstrating the potential of this instrument for clinical applications.

## 1. Introduction

Non-invasive cellular-scale imaging of the cornea is a valuable tool for disease diagnostics, management, and monitoring. In clinical practice, it is often used to distinguish forms of microbial keratitis *in situ*, when corneal biopsy is either infeasible or fails to yield a diagnosis [1]. It is used routinely to examine the endothelium for evidence of structural change or dysfunction, which can cause corneal edema and concomitant vision impairment [2]. Cellular-scale corneal nerve imaging has also been suggested as a means to monitor systemic disease, such as diabetes mellitus, or recovery from refractive surgery [3].

Currently, the established microscopic clinical imaging methods are specular microscopy (SM) [4] and *in vivo* confocal microscopy (IVCM) [5]. Whereas SM is restricted to the endothelium, IVCM is able to produce high contrast images throughout the full thickness of the human cornea and resolve nerves and cells in 3D [6]. A caveat is that IVCM is usually performed in contact with the cornea, meaning the objective lens (or protective cap) touches the cornea during the examination. Topical anesthetic must be administered prior to imaging. With an experienced technician, contact operation is safe and straightforward. Nevertheless, there are many subjects with phobias that will not tolerate contact operation. Moreover, for routine screening purposes where speed is critical, non-contact methods are highly desired.

Much of the recent progress in non-contact *in vivo* corneal imaging has involved various flavors of optical coherent tomography (OCT) [7, 8], largely due to its remarkable depth selectivity. Chen et al. described a micro-OCT system capable of cross-sectional swine cornea imaging [9]. However, the A-line rate was not fast enough to provide useful *en face* images in the presence of motion. Mazlin and colleagues took a different approach and applied a parallelized full-field version of OCT (FF-OCT). They were able to acquire very large *en face* images in human corneas [10]. With faster A-line rates, Tan et al. later showed high-resolution imaging in 3D was feasible with spectral-domain OCT [11]. Recently, Gabor-domain optical coherence microscopy has also been successfully demonstrated *in vivo*, albeit only in anesthetized mice [12].

All the techniques mentioned so far are based on reflection, or, more precisely, backscattered light from corneal microstructures. Here we introduce an *in vivo* microscopy technique based instead on transmitted light. We call this approach retroillumination microscopy, in deference to the related but lower-resolution slit lamp technique [13]. The key idea is to use the ocular fundus as a diffuse back-reflector, thereby folding the light path of a widefield transmission microscope into one which requires access to only one side of the sample (either the cornea or crystalline lens). To maximize back-reflection, we use near-infrared (NIR) light, which is weakly absorbed in the fundus and virtually undetectable to the subject. Additionally, we implement asymmetric illumination, a well-established method for enhancing intrinsic phase-gradient contrast [14–16]. Our method is non-contact and produces images with high lateral resolution, comparable to state-of-the-art IVCM systems, and across a large field in the cornea (1-mm diagonal). A strength of the system is its instrumentational simplicity, making it a promising candidate for disease screening or global-health applications.

The purpose of this report is to describe the retroillumination microscope design in detail. We also present representative images of the cornea and lens obtained from healthy volunteer subjects.

## 2. Device description

### 2.1. Hardware

A schematic for the retroillumination microscope is given in Fig. 1 and cross-sectional illustrations of the illumination beam at the cornea and fundus are given in Fig. 2. The subject’s head is placed on a chin rest, while their gaze is stabilized with an external fixation target. The microscope is mounted on translation stages for alignment with the eye.

**Fig. 1.**
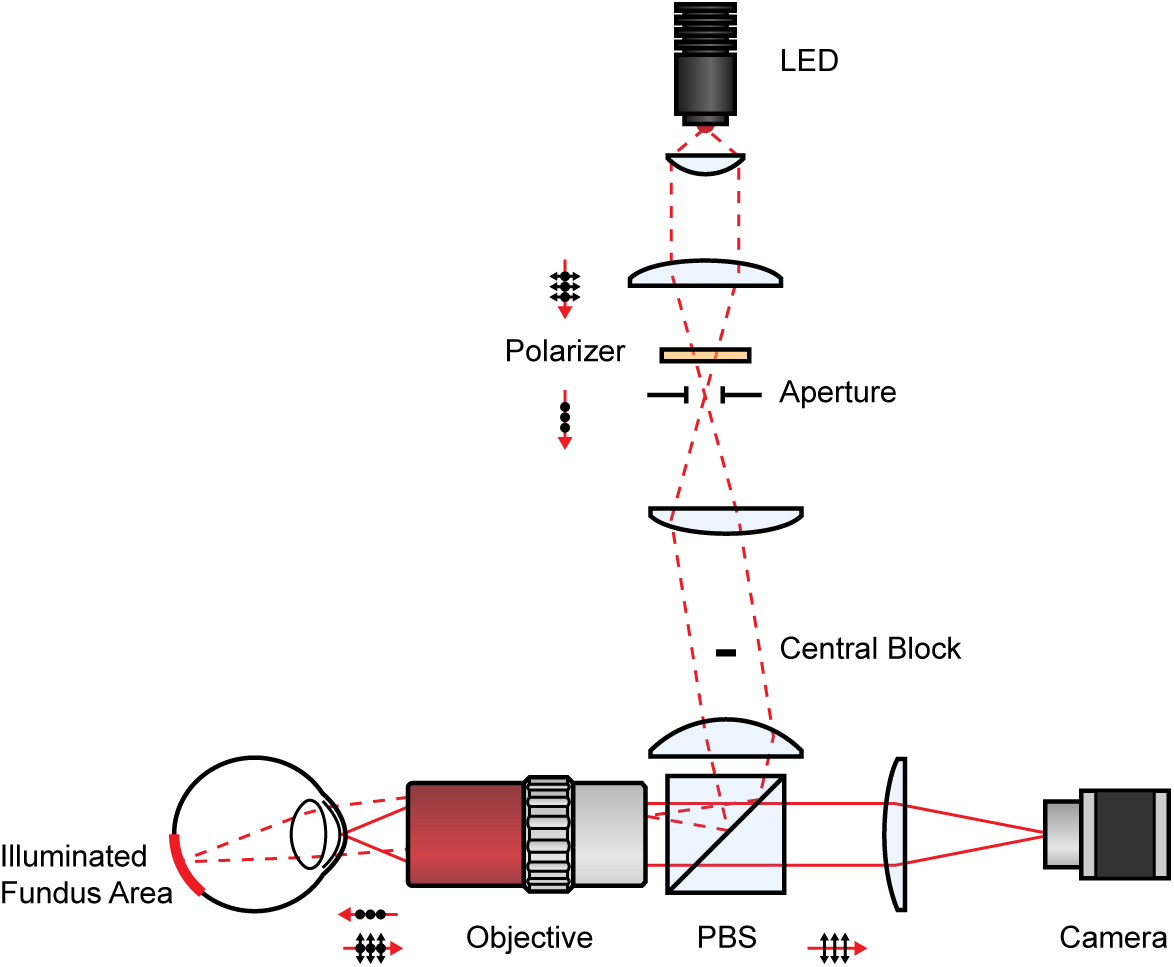
System optical diagram. PBS: Polarizing beamsplitter

**Fig. 2.**
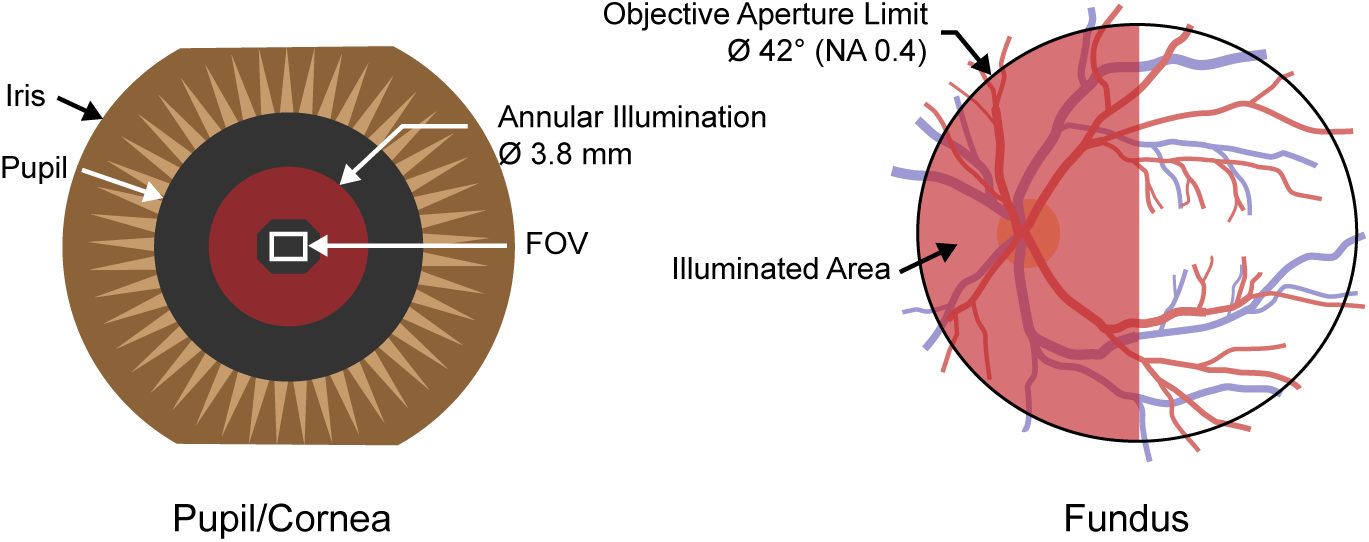
Illumination in cross-section at the pupil/cornea (left) and at the fundus (right).

Illumination consists of a high power NIR LED (LZ1-10R602, Osram; 850 nm center wavelength), which is offset and magnified to span a semi-circular aperture and subsequently relayed through a beamsplitter to the back focal plane of a long working distance objective lens (MPlanApoNIR 20X/0.4, Mitutoyo). The diameter of the aperture is adjusted such that its image half fills the objective’s back aperture. The objective lens then projects the semi-circular LED image onto the fundus with a visual angle of about 42° (right side of Fig. 2). The beam diameter at the pupil is about 3.8 mm (Fig. 2, left). The objective lens working distance is 20 mm, which allows the subject to freely blink.

We assume the fundus acts as a spatially uniform diffuse reflector. Features such as retinal vessels and the optic nerve head exhibit rather weak contrast with NIR illumination [17] and hence can be ignored. Reflected light from the fundus obliquely transilluminates (i.e. retroilluminates) the anterior segment. Rays clearing the iris are collected by the same objective and magnified with a compound tube lens onto the sensor of a machine vision camera (acA2000-340kmNIR, Basler), running at 348 frames/sec. We chose this speed based on speeds used successfully in corneal FF-OCT [10]. We used two off-the-shelf achromats (AC508-300-B and AC508-150-B, Thorlabs) spaced by about 2 mm to achieve the desired tube lens power while avoiding additional aberration.

The system described thus far is susceptible to direct backreflections from the objective and to a lesser extent, the anterior segment interfaces, which can severely impair image contrast. To mitigate direct backreflections, we use cross-polarized detection and exploit the fact that multiple scattering in the fundus largely depolarizes the retro-scattered illumination. Specifically, the LED light is linearly polarized (LPVIS100-MP2, Thorlabs) and combined with a polarizing beamsplitter cube (PBS252, Thorlabs).

In the illumination path, we also insert a small central block (mounted on a transparent window) conjugate to the cornea in order to darken the microscopy field of view (FOV) and reduce depolarized stray light. The block is responsible for the octagon-shaped hole in the incident illumination on the left in Fig. 2. Although this configuration resembles an ophthalmoscope configuration, with its spatially segmented input and output beams, we emphasize that the retroillumination microscope is focused on the anterior segment FOV, not the fundus.

In practice, it is difficult to align the microscope to the eye using only the high-resolution FOV. We obtain a wider field for alignment by using an auxiliary camera to image the subject’s pupil with ambient light. Specifically, a motorized flip mirror is inserted in the illumination path just after the aperture (omitted from Fig. 1 for clarity), which temporarily redirects an approximately 7 mm diameter low-resolution image of the pupil onto a free-running machine vision camera (DCC3240N equipped with camera lens MVL6WA, Thorlabs). We use the edges of the subject’s iris to then center the microscope prior to microscopic imaging. This alignment procedure increases operator repeatability. After locating the desired structures, we usually capture sets of 1024 frames (3 sec video).

### 2.2. Resolution and field of view

In the asymmetric retroillumination microscope, and indeed all microscopes based on asymmetric illumination, the phase-to-intensity point spread function is no longer an Airy pattern, making it difficult to ascribe a resolution based on, for instance, the Rayleigh criterion. Instead we report spatial frequency bandwidth as a surrogate for resolution. For 850 nm light and 0.4 NA (illumination and imaging), the maximum spatial frequency is laterally 940 mm^-1^ (1.1 µm period) and axially 98 mm^-1^ (10 µm period), in air. The camera produced images of dimensions 1540 x 1088 pixels. This corresponds to a field of view (FOV) of about 820 x 580 µm (or 1 mm diagonal) in the cornea.

### 2.3. Light levels and safety

The total incident light power on the cornea is about 50-100 mW. This light level (at 850 nm) does not cause significant pupil contraction and is just barely visible to the subject. Because the power is distributed over an area, the corneal and retinal irradiances are below the limits for non-hazardous Group 1 devices in the latest ophthalmic safety standard, the ANSI Z80.36-2016.

## 3. Image processing

Even with asymmetric retroillumination, intensity contrast at the image plane is low (<5%). Two major sources of noise restrict useful post-hoc expansion of this range: photoresponsive non-uniformity (PRNU) and shot noise. Note that offset noise is corrected on chip with correlated double sampling. Read and quantization noise are negligible. PRNU results from varying pixel gain and is easily corrected by dividing each raw frame by a calibration frame. We use a 256 frame average of uniform intensity, near the expected signal level, as the calibration frame.

Averaging several frames (equivalent to integrating more photons) reduces shot noise, but increases susceptibility to motion blur. Nevertheless, we could still average several frames by registering frames prior to averaging. Registration was performed with standard FFT-based phase-correlation methods [18]. We found that we can usually average at least 9 frames before axial motion decorrelates the FOV.

Additionally, we remove any slowly-varying illumination gradients by dividing the image by a Gaussian-filtered version of itself (σ = 24 pixels or 13 µm in the cornea). Processing is performed efficiently in a consumer GPU and a cropped version of the images is displayed in real time at approximately 20 Hz. Following acquisition, we perform the same processing on the full FOV and for each frame in the stack.

## 4. Example images

### 4.1. Subjects

We imaged the left eye of 3 healthy volunteers ranging in age from 26 to 57 and with varying fundus pigmentation. For each subject, informed consent was obtained prior to imaging. The research was approved by the Boston University Institutional Review Board and conformed to the principles stated in the Declaration of Helsinki.

### 4.2. Epithelium

The subbasal nerve plexus (SBP) was the most visible structure observed in the epithelium. The SBP is a highly-branched network of sensory nerve fibers located in a narrow plane between the basal epithelial cell layer and Bowman’s layer. Figure 3(A) shows the SBP across nearly the entire FOV close to the corneal apex of a 28-year-old male. The large dark spots are out-of-focus aggregates of either mucin or shedded epithelial cell debris on the superficial cornea, anterior to the focal plane. These spots appear dark because the aggregates scatter light outside the collection aperture resulting in a decrease in local image intensity. The aggregates usually shift over time and move rapidly following a blink.

**Fig. 3.**
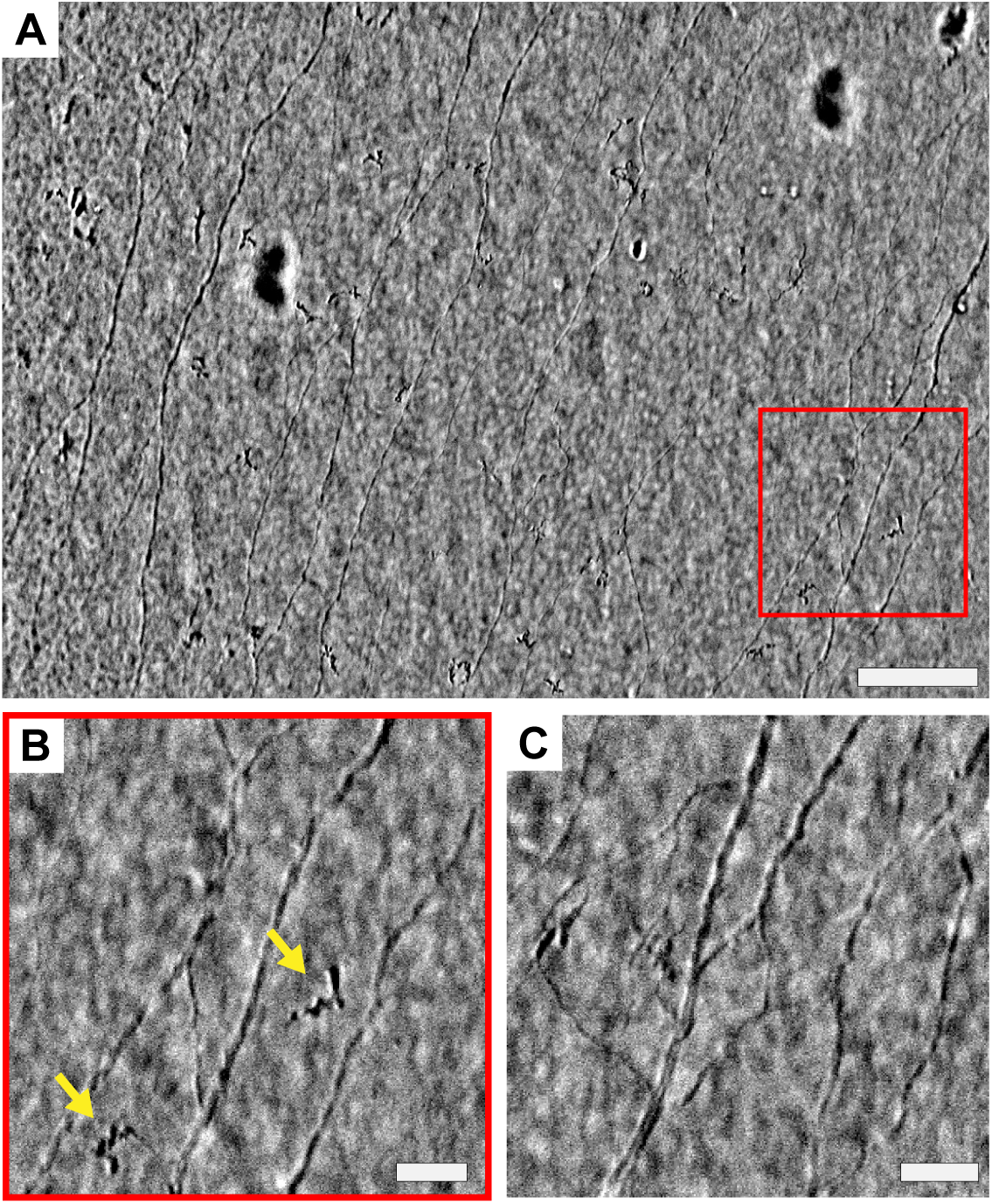
Subbasal nerve plexus of a 28-year-old male near the corneal apex. (A) Full FOV (820 x 580 µm) and (B) indicated area in detail. Arrows indicate suspected Langerhans cells or intraepithelial fibers. See Visualization 1 for a full FOV video excerpt. (C) Another area seen later in the image sequence showing complex anastamoses. Scale bars: (A) 100 µm, (B) & (C) 25 µm.

Enlarged areas are given in Figure 3(B) & (C) showing the complex SBP branching and variation of fiber diameter. We also repeatedly saw globular structures (blue arrows) near or just anterior to the SBP plane of focus. These structures are more discernible in the video. They may be Langerhans cells (resident macrophages) or intraepithelial nerve fibers.

Other periepithelial features are shown in Figure 4. The tear layer has a punctate appearance (Fig. 4(A)) with sparse highly-scattering aggregates. The edges of squamous and wing epithelial cells are occasionally visible (Fig. 4(B)), but have low contrast and are therefore difficult to distinguish. The basal epithelial cell mosaic (Fig. 4(C)) is always visible with positive or negative contrast depending on the relative position in the focal plane. In one subject, we saw numerous dendritic cells with clearly resolvable cell bodies and processes (Fig. 4(D)). We were unable to detect any discernible features in Bowman’s layer.

**Fig. 4.**
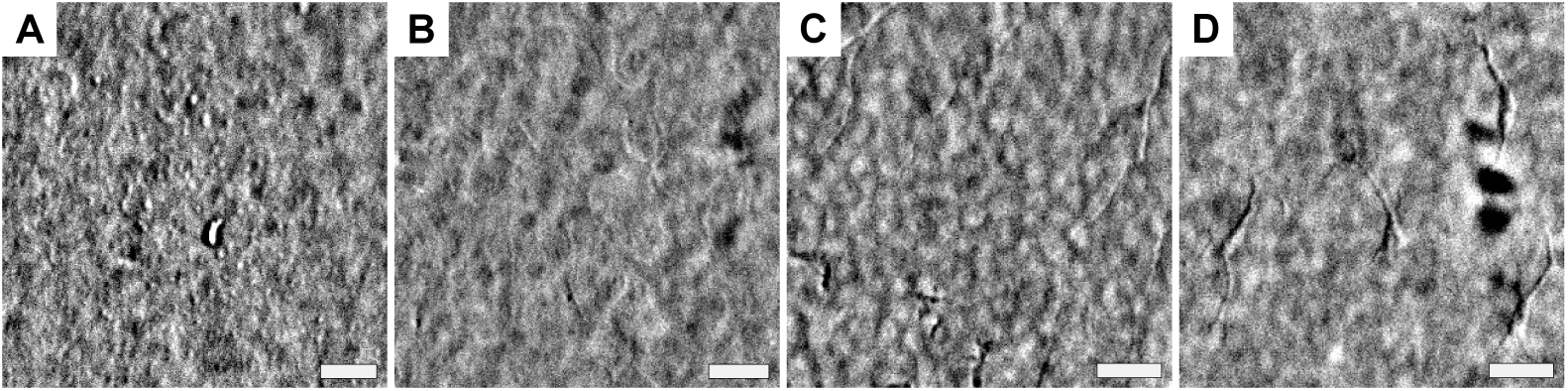
Other epithelial structures. (A) The tear layer of a 26-year-old female. (B) Superficial and (C) basal epithelial cells in a 28-year-old male. (D) Dendritic immune cells in a 57-year-old male. Scale bars: 25μm.

### 4.3. Stroma

Within the stroma, we observed large branching nerves. In contrast to subbasal nerves which are largely confined to a plane, stromal nerves are distributed throughout the stromal volume. Hence, it was challenging to obtain images where large portions of the nerve were in focus. Figure 5 shows a few stromal nerve segments. Panels A and B are cropped views of the same stromal nerve trunk but at slightly different focal planes. Arrows indicate the common branch point.

**Fig. 5.**
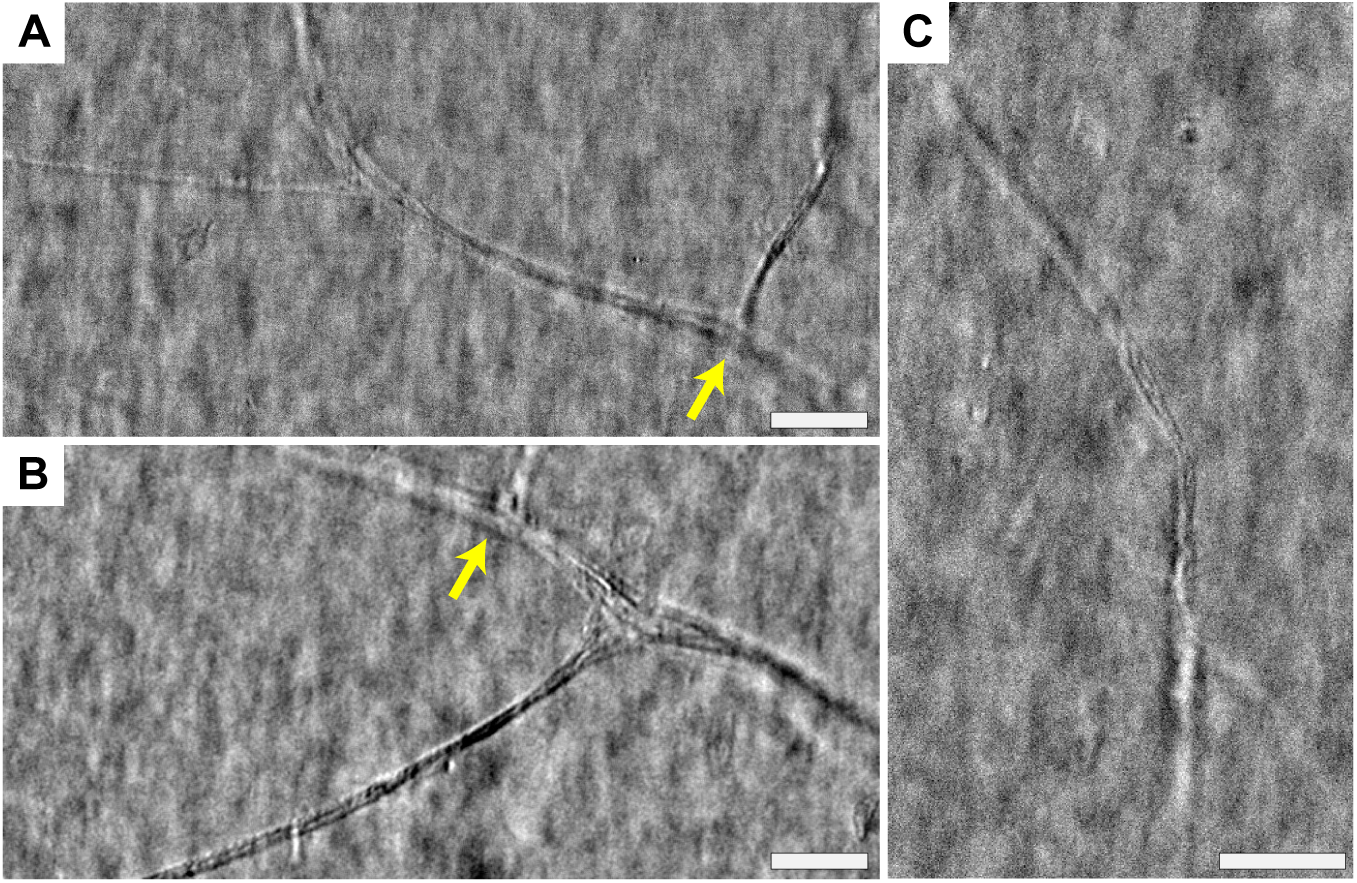
Stromal nerves in a 28-year-old male. (A) Stromal nerve segment just anterior to view shown in (B). Arrows indicate the same branch point. (C) Stromal nerve in focus near the center and out of focus at the top and bottom. Scale bars: 50 µm.

Keratocyte nuclei were notably absent from our retroillumination images. This was surprising based on their density in the stroma and high contrast in IVCM [5]. In video sequences, we occasionally observed distinct structures about the size of a cell, but in much lower abundance than expected for normal keratocytes. We also did not see any indication of the subepithelial nerve plexus, however we confined most of our imaging the central cornea where the subepithelial plexus may simply be absent [19].

### 4.4. Endothelium

The endothelium, a monolayer of cells that coats the posterior surface of the cornea, was readily visible with retroillumination. A widefield 1-mm diameter view is shown in Figure 6. Interestingly, it appears that endothelial cell nuclei, and not cell edges, exhibit the best contrast. Similar to basal epithelial cells, when endothelial cell nuclei are above (i.e. anterior to) the focal plane, they produce positive contrast and conversely, when they are below the focal plane, they produce negative contrast. The curvature of the endothelium is also evident in Figure 6. Just anterior the endothelium, at the approximate location of Deschemet’s membrane, we also repeatedly saw small high contrast spots. These spots were much smaller than the size of the nearby endothelial cells.

**Fig. 6.**
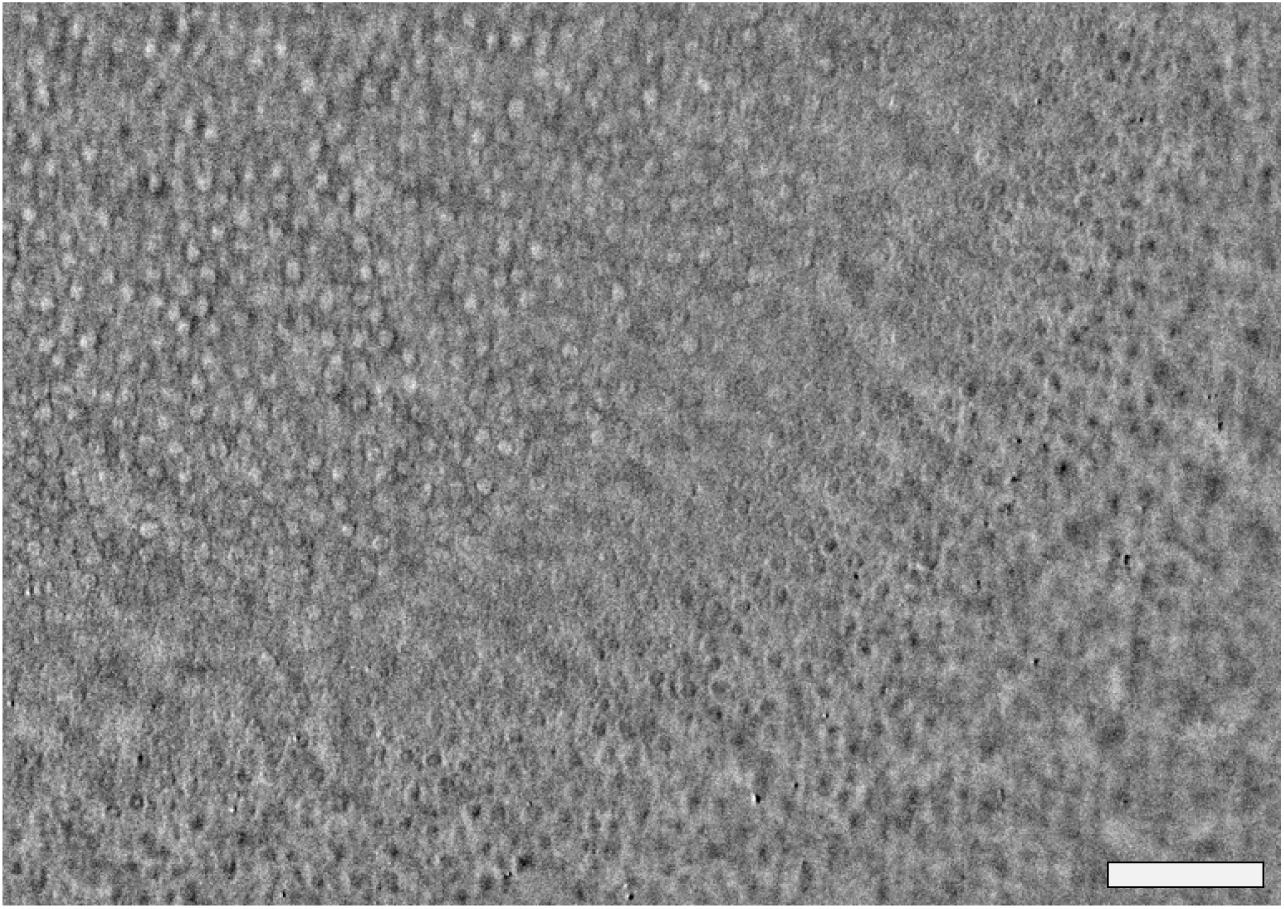
Endothelium of a 26-year-old female. Bright/dark circles are nuclei anterior or posterior the focal plane (see Visualization 2 for through-focus). The curved zone of null contrast suggests that the corneal apex is located near the top left corner of the image. Scale bar: 100 µm.

### 4.5. Crystalline lens

With a 20 mm working distance objective lens we can easily adjust the system to image the crystalline lens, which begins about 3.3 mm behind the air-cornea interface. However, ray tracing software indicated that strong spherical aberration, from the air-cornea refractive index mismatch, impedes clear imaging. To reduce sensitivity to spherical aberration, we lowered the effective imaging NA by relaying the objective’s back focal plane to an external iris prior to focusing on the camera. Passage through the iris reduced the imaging NA down to 0.2. Despite the lower resolution, both the lens epithelium and anterior lens fibers were discernible. Example images from a 28-year-old male are shown in Figure 7.

**Fig. 7.**
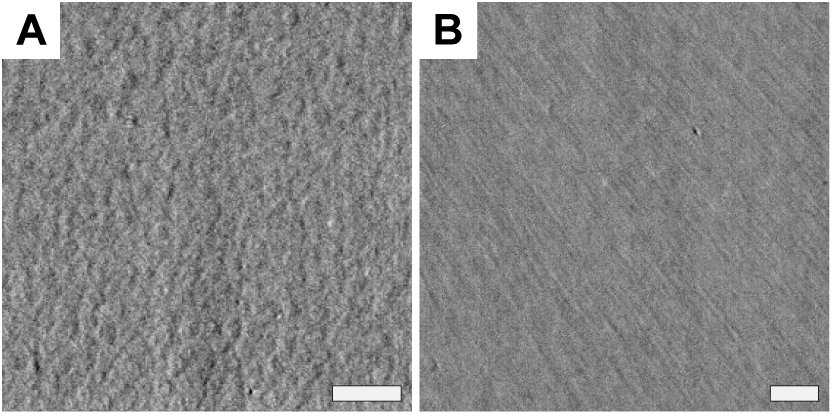
Crystalline lens (A) epithelium and (B) anterior fibers just below the epithelium. Scale bars 50 µm

## 5. Discussion

We have described a new *in vivo* corneal and lenticular imaging method, which we call retroillumination microscopy. The technique is non-contact and produces images with high lateral resolution (1-2 µm), comparable to state-of-the-art IVCM.

Unlike most other *in vivo* eye imaging techniques, retroillumination microscopy is based on transmitted light. This difference has a fundamental impact on obtainable image contrast [8, 20]. In order for light reflection, or more precisely, backscattering to occur, the sample must present an abrupt change in refractive index. This could either be an interface (specular reflection) or a clump of scattering structures each smaller than the wavelength of incident light (Rayleigh-like scattering). On the contrary, transmission microscopy is sensitive to forward-scattered light, such as that primarily generated by larger structures, for example cell bodies or nuclei. A clear example is the different appearance of the corneal endothelium, which in reflection contrast usually appears as a hyper-reflective interface with dark paths delineating cell borders [4]. In transmission contrast, it is the cell nuclei that are most apparent, while cell edges are undetectable (see Fig. 6). As an aside, the adaptive optics ophthalmoscopy community has already recognized the utility of forward scattered light as a method to enhance contrast of blood flow [21], photoreceptor cone inner segments [22], and retinal ganglion cells [23]. There are likely many other corneal features where transmission contrast can contribute complimentary information.

Our method bears resemblance to the well-known slit lamp technique known as retroillumination [13] (n.b. the naming similarity is intentional). Both techniques reflect light off posterior structures in order to back-illuminate the lens and cornea. However, unlike the slit lamp biomicroscope, which features separate, non-overlapping illumination and imaging paths, our system is designed around a single objective lens. With this configuration we are free to use a much higher collection angle (NA) in the imaging path without physically obstructing the illumination. Thus our single lens configuration provides higher optical resolution than that obtainable with standard slit lamps. Similarly, our design enables unimpeded illumination of large fundus areas. We use this freedom to implement asymmetric illumination, a well-established method to enhance phase-gradient contrast [14].

Transmission imaging also avoids superficial sample reflections, such as the prominent corneal anterior surface reflection. Excess background from this reflection can easily dominate intracorneal backscattering. Hence, high optical sectioning strength (e.g. confocal filtering or coherence gating) is normally required to diminish its effect, which in turn increases system complexity. In the absence of this reflection, we are able to form useful *en face* images across a large, 1-mm diagonal FOV with little more than a widefield microscope made of readily available off-the-shelf components.

Retroillumination provides excellent contrast of corneal nerves, particularly the subbasal plexus, which is recognized as a potential biomarker for diabetic peripheral neuropathy [24]. Combined with the large FOV (3X larger area than current IVCM) and non-contact operation, retroillumination microscopy may be a useful tool for monitoring diabetic neuropathy or other ocular diseases affecting corneal nerves.

